# Apicosome: newly identified cell-type-specific organelle in mouse cochlear and vestibular hair cells

**DOI:** 10.1101/2022.07.20.500729

**Authors:** Xiaofen Li, Qirui Zhao, Xiaojie Yu, Wenhan Cao, Yingyi Zhang, Wanying Feng, Liwen Jiang, David Z. He, Robert Z. Qi, Pingbo Huang

**Affiliations:** Division of Life Science, Hong Kong University of Science and Technology, Hong Kong, China; Department of Chemical and Biological Engineering, Hong Kong University of Science and Technology, Hong Kong, China; State Key Laboratory of Molecular Neuroscience, Hong Kong University of Science and Technology, Hong Kong, China; HKUST Shenzhen Research Institute, Hong Kong University of Science and Technology, Hong Kong, China; Hong Kong Branch of Guangdong Southern Marine Science and Engineering Laboratory (Guangzhou), Hong Kong University of Science and Technology, Hong Kong, China; Biological Cryo-EM Center, Hong Kong University of Science and Technology, Hong Kong, China; Office of the Vice-President for Research and Development, Hong Kong University of Science and Technology, Hong Kong, China; School of Life Science, The Chinese University of Hong Kong, Hong Kong, China; Department of Biomedical Sciences, Creighton University School of Medicine, Omaha, NE, USA

## Abstract

Cochlear and vestibular hair cells in the inner ear are highly specialized sensory receptors for sound waves and acceleration of body movements; these cells can perform their specialized functions because of their distinctive morphology and some unique organelles that they harbor. Here, we report a serendipitous identification in the mouse of a hair-cell-specific organelle, which we name “apicosome.” The apicosome was recognized by anti-FLRT1 antibodies but contains no FLRT1, and the organelle presents several distinctive characteristics: (1) the apicosome typically appears as a single entity (∼500 nm in diameter), but occasionally as two entities, in hair cells; (2) it first appears in the subapical region at the neural side at embryonic day (E) 17–18 in cochlear hair cells, subsequently descends to the perinuclear region during the first postnatal week, and completely disappears around postnatal day (P) 10; (3) in vestibular hair cells, it can be detected in the subapical region of neonatal (P3) cells and persists in adult hair cells although it becomes smaller and more distant from the subapical region; (4) the timing of apicosome translocation and disappearance during development is correlated in kinocilium maintenance; (5) the organelle is potentially associated with microtubules; and (6) the appearance of the apicosome is irregular in supernumerary hair cells and this is likely linked to anomalous lateral inhibition. Thus, our study identifies a previously undescribed organelle in sensory hair cells and lays the foundation for further characterization of this specialized structure potentially linked to hair-cell development and morphogenesis.

## INTRODUCTION

A cellular organelle is a specialized subunit responsible for a distinct function in the cell. Organelles can be either membrane-bound, such as the nucleus and mitochondrion, or non-membranous, such as the ribosome and centrosome. Whereas several organelles are present in most cell types, other organelles are highly specific to certain cell types, such as synaptic vesicles in neurons, melanosomes in melanocytes, Weibel-Palade bodies in endothelial cells, and chloroplasts in plant and algal cells. These cell-type-specific organelles are formed to serve cell-specific biological functions.

Sensory hair cells in the inner ear of vertebrates, including cochlear and vestibular hair cells, also harbor cell-specific organelles and structures required for performing their highly specialized functions—to detect sound waves and acceleration of body movements. For instance, the stereociliary bundle, which is composed of actin-based cilia (ranging in number from tens to over one hundred) displaying a staircase arrangement, is a sensory organelle through which mechanical stimuli are detected by the hair-cell transduction machinery—the mechanotransduction (MT) complex. Another example is the microtubule-based kinocilium present in vestibular hair cells and immature cochlear hair cells, but not in adult mammalian cochlear hair cells; the kinocilium inserts into the cell body at the fonticulus located at the abneural side of postnatal hair cells and is associated with the basal body. The kinocilium is considered to play a critical role in hair-bundle morphogenesis. Other hair-cell-specific organelles and structures include the cuticular plate, synaptic ribbon, striated organelle (SO, or the Friedmann body) (Pollock and McDermott Jr, 2015), and Hensen’s body (HB) (Lim, 1986; Mammano et al., 1999).

Here, we report the discovery of an organelle specific to cochlear and vestibular hair cells. We coined the term “apicosome” for this previously undescribed organelle, which exhibits a distinctive shape, size, translocation dynamics, and developmental profile. We also discuss and suggest here the potential function of the apicosome. Our work lays the foundation for further characterization of the apicosome, an organelle potentially linked to hair-cell development and morphogenesis.

## MATERIALS AND METHODS

### Antibodies and phalloidin

The following primary antibodies were used in this study: two rabbit polyclonal antibodies against human fibronectin leucine-rich transmembrane protein 1 (FLRT1) (ab97825 and ab103839; Abcam, Cambridge, UK), rabbit polyclonal anti-human FLRT2 (ab154023; Abcam), mouse monoclonal anti-human FLRT2 (MAB2877; R&D Systems, Minneapolis, MN, USA), goat polyclonal anti-human FLRT3 (AF2795; R&D Systems), mouse monoclonal anti-centrin (04-1624; EMD Millipore, Darmstadt, Germany), and mouse monoclonal anti-HA (901502; BioLegend, San Diego, CA, USA). The secondary antibodies used were Alexa Fluor 488-conjugated goat anti-rabbit IgG (A27034; Thermo Fisher Scientific, Waltham, MA, USA), goat anti-mouse IgG (A28175; Thermo Fisher Scientific), donkey anti-goat IgG (ab150129; Abcam); Alexa Fluor 647-conjugated goat anti-mouse IgG (ab150115; Abcam); and biotinylated goat anti-rabbit IgG (65-6140; Thermo Fisher Scientific). For labeling F-actin, we used TRITC-conjugated phalloidin (P1951; Sigma-Aldrich, St. Louis, MO, USA) or Alexa Fluor 647-conjugated phalloidin (A22287; Thermo Fisher Scientific).

### *Flrt1*^*-/-*^ mice and genotyping

FLRT1-knockout (KO; *Flrt1*^*-/-*^) mouse sperm was purchased from MMRRC (B6.129S5-*Flrt1*^*tm1Lex*^/Mmucd; RRID: MMRRC_032313-UCD) to generate KO mice with the C57BL/6 background. All animal procedures were approved by the University Committee on Research Practices at the Hong Kong University of Science and Technology (Ethics Protocol: 2018001).

*Flrt1*^*-/-*^ mice were genotyped using these primers: 5ʹ-GCACCACACGAGGCTACCG-3ʹ (sense) and 5ʹ-ACATCTCCAACAATGCTGAATCCC-3ʹ (antisense), designed to produce a 271-bp wild-type (WT) band; and 5ʹ-GCAGCGCATCGCCTTCTATC-3ʹ (sense) and the WT antisense primer, to produce a 371-bp KO band. Amplification reactions were performed using Taq DNA polymerase at 61°C annealing temperature.

### Cell culture and transfection

Cells were cultured and transfected according to our published procedures (Hu et al., 2017). HEK293T cells (RRID: CVCL_1926) and COS7 cells (RRID: CVCL_0224) were obtained from American Type Culture Collection (ATCC) (Manassas, VA, USA); the cells were presumably authenticated by ATCC and were not further authenticated in this study. The cell lines, which routinely tested negative for mycoplasma contamination, were maintained in DMEM supplemented with 10% fetal bovine serum (FBS) and 100 μg/mL ampicillin in an atmosphere of 95% air and 5% CO_2_ at 37°C. All transfections were performed using polyethylenimine (Polysciences, Inc., Warrington, PA, USA). Cells were transiently transfected with pcDNA3-Flrt1-HA or pcDNA3-Flrt2-HA, and, after culturing for 48 h, were subject to western blotting or immunostaining.

### Cochlear culture and treatments

Cochlear culture was performed following a published protocol (Xiong et al., 2014). Briefly, the organ of Corti was dissected from postnatal day (P) 2 mice, attached on the bottom of a 35-mm culture dish, and incubated in DMEM/F-12 medium containing 1% (v/v) FBS and 1.5 μg/mL ampicillin at 37°C overnight before treatment. Cochlear cultures were treated for 6 h with various reagents (all dissolved in culture medium): 0.1% (v/v) DMSO (276855; Sigma-Aldrich), 0.5 mM BAPTA-AM (A1076; Sigma-Aldrich), 1 μM thapsigargin (T9033; Sigma-Aldrich), 0.1 mM dihydrostreptomycin (DHS) (D1954000; Sigma-Aldrich), 0.1 mM JW67 (SML0324; Sigma-Aldrich), or 100 ng/mL epidermal growth factor (EGF) (PHG0311; Thermo Fisher Scientific).

To block Notch signaling, DAPT, a γ-secretase inhibitor, was used as previously described (Li et al., 2018). The cultures prepared from P0 WT mice were treated with 5 µM DAPT (CAS208255-80-5; Santa Cruz Biotechnology, Dallas, TX, USA) or 0.05% (v/v) DMSO (vehicle) for 48 h; the cultures were then washed and incubated for additional 48 h in fresh culture media. The samples were fixed, immunostained, and imaged. At least three cochleae from WT mice were used for each experimental condition.

### Immunohistochemical assays

Cochlear tissues and COS7 cells were immunostained following our published procedures (Li et al., 2019). Briefly, tissues were fixed with 4% (w/v) paraformaldehyde (PFA) or ice-cold methanol, immersed in HBSS containing 0.1 mM CaCl_2_, 4% (w/v) bovine serum albumin (BSA), and 0.5% (v/v) Triton X-100 for blocking and permeabilization, and then incubated with primary antibodies (1:200 anti-FLRT1, anti-FLRT2, or anti-FLRT3) at room temperature for 4 h. Subsequently, the samples were washed thrice (10 min each) with HBSS containing 0.1 mM CaCl_2_ and incubated with Alexa Fluor 488-conjugated 2^nd^ antibodies (1:200) and Alexa Fluor 647-conjugated phalloidin (1:40) at room temperature for 2 h. After washing thrice (10 min each) with HBSS containing 0.1 mM CaCl_2_, the samples were mounted using ProLong Gold Antifade Mountant (P36930; Thermo Fisher Scientific).

In the case of COS7 cells, at 48 h after transfection with FLRT1-HA or FLRT2-HA, the cells were fixed with 4% PFA for 1 h, washed once with PBS, blocked and permeabilized with PBS containing 4% BSA and 0.5% Triton X-100 for 1 h, and then incubated with primary antibodies (1:200) at room temperature for 4 h. After washing thrice (10 min each) with PBS, the cells were incubated with Alexa Fluor 488-conjugated goat anti-rabbit IgG (1:200) and Alexa Fluor 647-conjugated goat anti-mouse IgG (1:200) plus TRITC-conjugated phalloidin (1:400) at room temperature for 2 h. Lastly, the cells were washed thrice (10 min each) with PBS and mounted for examination under a confocal microscope.

All confocal microscopy was performed using Leica TCS Sp5 or Leica TCS Sp8 confocal microscopes (Leica Microsystems, Buffalo Grove, IL, USA) (objectives: 100× for whole-mount cochleae; 63× for whole-mount vestibular tissues and COS7 cells; 1.4 NA). Leica LAS-AF and LAS-X imaging software and Fiji software (https://fiji.sc) were used for image acquisition and analysis.

### Transmission electron microscopy (TEM) and immunogold labeling

Cochleae collected from P2 WT mice were fixed with 4% (w/v) PFA in 0.1 M sodium phosphate buffer (PB, pH 7.4) at room temperature for 2 h, and then kept at 4°C for 12 h; the cochleae were punched with small holes in apex and base areas for better perfusion. After washed thrice with 0.1 M PB, the organs of Corti were then dissected from the fixed cochleae. For post fixation, the organs of Corti were treated with 1% (w/v) osmium tetroxide (OsO_4_) in 0.1 M PB for 20 min at room temperature and washed with 0.1 M PB for at least 5 times (OsO_4_ fixation should be skipped if the samples were subject to immunolabeling). The organs of Corti were dehydrated in a graded series of acetone solutions (50%, 70%, 90%, 100%, and dry 100%). Then, the dehydrated organs of Corti were infiltrated in a mixture of acetone and Embed 812 resin (14120; Electron Microscopy Sciences), and polymerized in fresh pure resin at 60°C for 24 h.

The organs of Corti embedded in Embed 812 resin were sectioned with a histo-diamond knife (Diatome) on the ultramicrotome (Leica EM UC7, Leica Microsystems). Ultra-thin sections (80 nm in thickness) were made and collected on 3 mm-diameter nickel grids.

If immunolabeling was not required, the grids were only stained with 2% uranyl acetate for 10 min and 1% lead citrate for 5 min. While for immunogold labelling, grids were immersed in droplets of the following solutions sequentially within a humid chamber: tris buffered saline (TBS) to wash once; 10% (v/v) rabbit serum and 0.2% (v/v) Tween 20 in TBS for 30 min at room temperature to block nonspecific protein binding; anti-FLRT1 antibody (ab97825, 1:30) in 0.05 M TBS containing 1% (w/v) BSA (BSA-TBS) at 4°C overnight; 0.5% (v/v) Tween 20 in TBS (T20-TBS) for 5 times (5 min each); biotinylated goat anti-rabbit IgG (1:100) in BSA-TBS for 1.5 h at room temperature; T20-TBS twice (5 min each); BSA-TBS twice (5 min each); STP-gold (1:10) in BSA-TBS for 1 h at room temperature; T20-TBS twice (5 min each); distilled water thrice (5 min each). Grids containing sections as negative controls were incubated in BSA-TBS without the primary antibody. Then, the grids were lightly counterstained with 2% uranyl acetate for 10 min and 1% lead citrate for 2 min.

As for imaging, the EM grids were screened using Tecnai transmission electron microscope (Thermo Fisher Scientific), operated at 120 keV. Images were taken with Ceta camera with the desired calibrated magnifications at the Biological Cryo-EM Center of the Hong Kong University of Science and Technology.

### Auditory brainstem response (ABR)

ABR measurements were obtained using our published procedures (Li et al., 2019).

## RESULTS

When WT mouse cochleae were fixed with PFA and immunostained with an antibody against FLRT1 (ab97825, Abcam), the signal appeared predominantly as a single distinct, large spot in the peri-cuticular necklace at the neural (modiolar) side of each inner hair cell (IHC) and outer hair cell (OHC) at P0 (Fig. 1A, upper panels), and the spot showed a clearly planar and apicobasal polarized distribution. The spots detected in different hair cells were mostly uniform in size (Supplementary Video 1), and the apparent size variation in Fig. 1A probably reflects the variation in the longitudinal positions. In control experiments, the 2^nd^ antibody (Alexa Fluor 488-conjugated goat anti-rabbit IgG) per se did not recognize the structure (Fig. S3C). We coined the term “apicosome” to describe this newly identified structure because it initially localized in the subapical region of hair cells (see below and Fig. 6); the structure is hereafter referred to as the apicosome.

**Figure 1.**
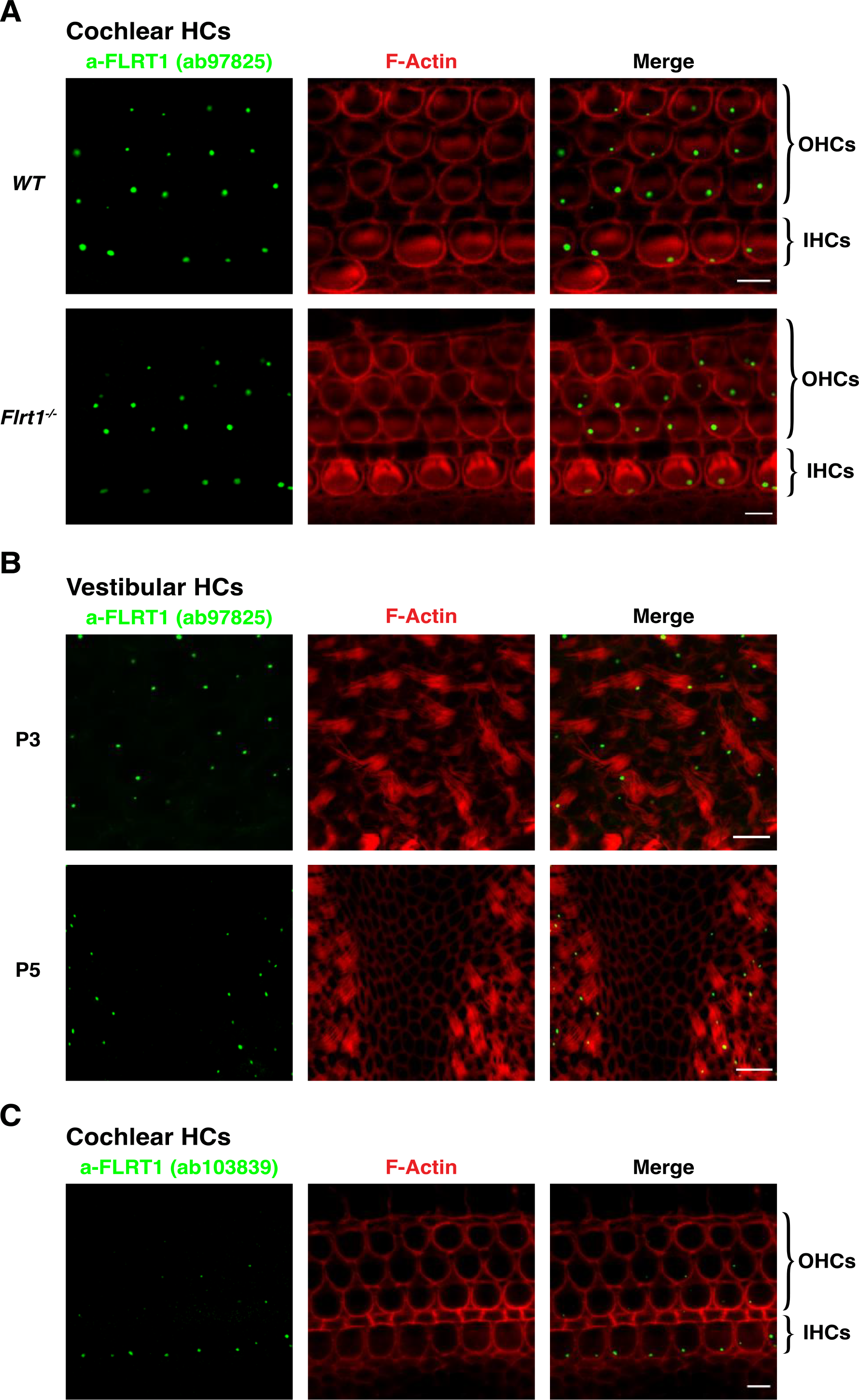
Hair-cell-specific spot-like structure recognized by anti-FLRT1 antibodies. (**A**) A cochlear hair-cell-specific structure, the apicosome, was recognized by an anti-FLRT1 antibody (ab97825, Abcam) in cochlear whole mounts from P0 WT (upper) and P1 *Flrt1*^*-/-*^ (lower) mice; the structure was absent in surrounding supporting cells. (**B**) The structure was also detected in vestibular hair cells from P3 (upper) and P5 (lower) WT mice and was again absent in supporting cells (middle region of lower panels). Because the apical plane of vestibular cells is not even and the hair bundles are long and floppy, the image of each channel was composed from stacks of images acquired in the same area of the tissue but at distinct focal planes within a depth of ∼15 μm in order to concurrently display the structure and the hair bundle in multiple hair cells. (**C**) The apicosome was also recognized (albeit weakly) by a different anti-FLRT1 antibody (ab103839, Abcam) in cochlear hair cells from P2 WT mice. Phalloidin staining shows F-actin (red). Scale bars: 5 μm (A, C) and 10 μm (B).

Interestingly, the antigen recognized by anti-FLRT1 in apicosomes is not FLRT1: the apicosome was also recognized by anti-FLRT1 in the hair cells of *Flrt1*^*-/-*^ mice (Fig. 1A, lower panels). Our immunostaining and immunoblotting results confirmed the specificity of the anti-FLRT1 antibody (ab97825, Abcam). The antibody staining closely colocalized with the HA-staining signal of FLRT1-HA expressed in cells (Fig. S1A), but the antibody weakly detected FLRT1 in western blotting and detected a strong nonspecific band of similar size (Fig. S1B). In another set of experiments, a different anti-FLRT1 antibody (ab103839, Abcam) (Fig. S1B) also detected the apicosome in WT hair cells, although the signal intensity was weaker (Fig. 1C). The antigens used for generating the two antibodies are aa 63–253 (ab97825) and aa 50150 (ab103839) of FLRT1, both of which are in the extracellular leucine-rich repeats of the protein. Our results suggest that the apicosome might contain a protein harboring a sequence similar to those of the two antigens, particularly the overlapping sequence: aa 63–150 of FLRT1 (see *Discussion*).

The apicosome was also detected in P3 and P5 WT vestibular hair cells beneath the apical surface (Fig. 1B), and the size and localization of the apicosome were highly consistent in all cochlear and vestibular hair cells. By contrast, the apicosome was not seen in the surrounding supporting cells (Fig. 1). These results indicate that the apicosome is a cochlear and vestibular hair-cell-specific organelle.

Next, to examine whether the apicosome is a microtubule-associated organelle, we used methanol instead of PFA to fix cochleae; this is because microtubules are fixed well by methanol, which denatures proteins but enables superior epitope exposure, whereas PFA generally preserves native structures. Interestingly, when cochleae were fixed with methanol, the apicosome appeared as a ring-shaped structure with an apparent diameter of ∼500 nm rather than as a concrete spot (Fig. 2A-C). These results suggest that the apicosome is a membrane-bound organelle and further that it might be associated with microtubules because the ring-like shape is detected when microtubule integrity is preserved. To observe the precise structure of the apicosome, we conducted transmission electron microscopy (TEM) with or without immunogold labelling on cochleae from P2 WT mice. In the non-labeled specimen, a round-shaped organelle clearly located at the neuronal side of subcuticular region in the hair cell, which can be easily distinguished from mitochondria or other conventional organelles around it (Fig. 2D, left). With FLRT1 immunogold labelling, the edge of this organelle were bound by the immunogold particles (Fig. 2D, right), indicating that the round-shaped structure is the apicosome that we identified in immunofluorescence experiments. The size and localization of the apicosome observed by TEM were consistent with those observed by confocal microscopy after immunofluorescence labeling. Thus, as verified by TEM, the apicosome is comparable in size (diameter) to other organelles such as mitochondria (0.5–1 μm), lysosomes (0.1–1.1 μm), and melanosomes (0.5 μm).

**Figure 2.**
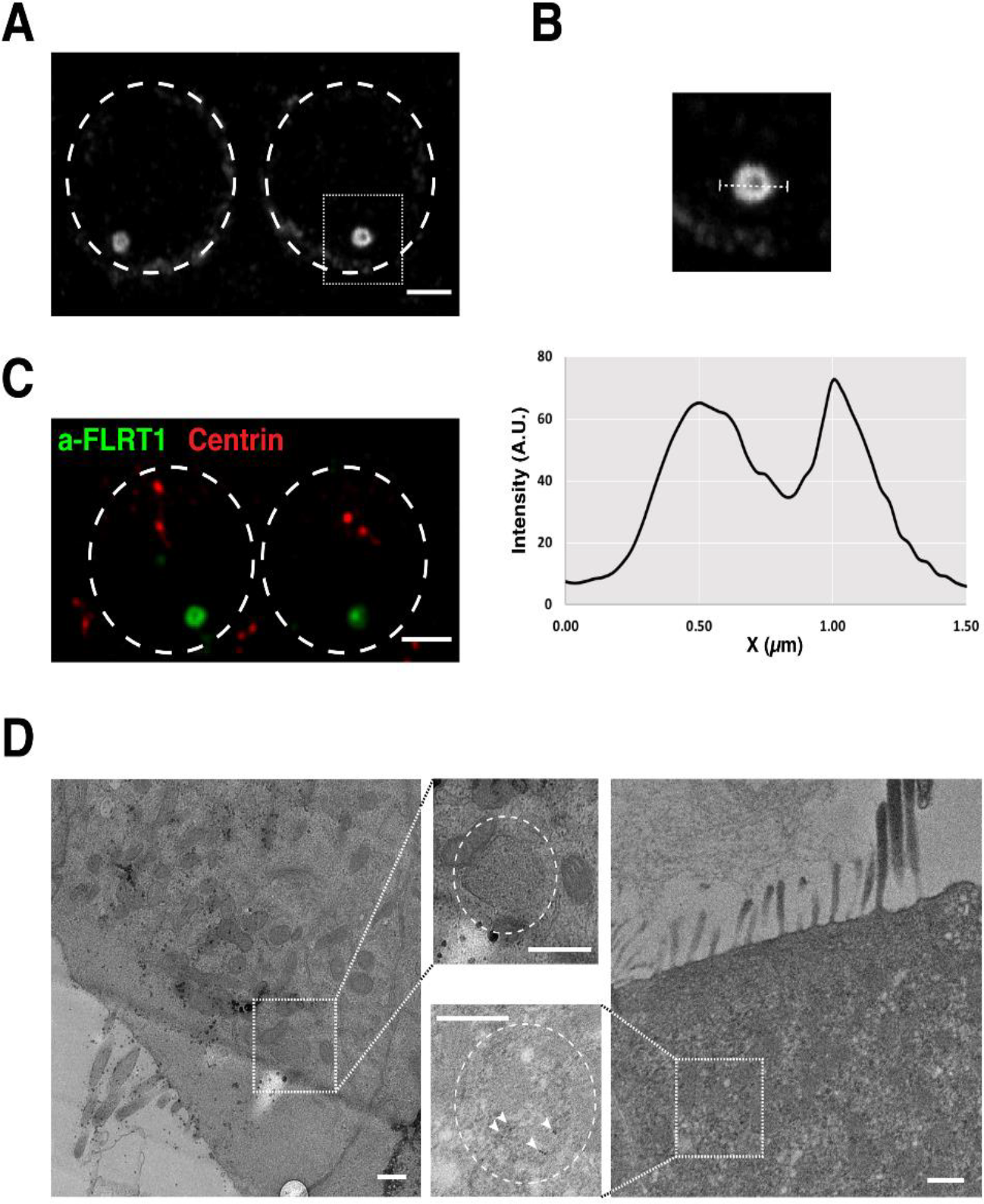
Apicosomes display a ring-like pattern with a diameter of ∼500 nm observed by confocal microscopy after methanol fixation and TEM. The annulus of the apicosome is shown (**A**) as a top view in P5 IHCs and (**C**) in P0 IHCs with co-staining for centrin (red). (**B**) Image: magnification of dotted box in (A); graph: fluorescence-intensity profile of the apicosome over the 1.5-μm-long dotted line; the distance of the two fluorescence-intensity peaks from the apicosome edge gives an estimation of the apicosome diameter (∼500 nm); A.U., arbitrary unit. In (C), the apicosome in the hair cell on the right is slightly out of focus, and centrin labeling in supporting cells is also visible. A round-shaped organelle with the diameter of ∼500 nm can be identified without (D, left) and with (D, right) FLRT1 immunogold labelling by TEM. The magnifications of the dotted boxes are shown in the middle. Dashed circles in (A) and (C): hair-cell boundaries. Dashed circles in middle (D): apicosomes. Arrowheads in (D): immunogold particles. Scale bars: 2 μm (A, C) and 500 nm (D).

The annulus observed here is reminiscent of what has been reported for recycling endosome clusters or vesicles surrounding centrosomes or non-centrosomal microtubule-organizing centers (ncMTOCs) (Sanchez and Feldman, 2017). In hair cells, the centrosome transforms into the basal body, which controls the formation and relocation of the kinocilium (Sienknecht, 2015), but the basal body localizes at the fonticulus in postnatal hair cells, i.e., on the opposite side of the apicosome (Fig. 6). We first suspected that the apicosome might be an organelle resembling the centrosome because (1) the apicosome appears as a single entity (or occasionally as two entities) in each hair cell; and (2) the centrosome is occasionally observed at the neural side of hair cells instead of at the opposite side like the basal body (Vranceanu et al., 2012). However, the apicosome did not colocalize with various centrosomal markers, such as centrin (Fig. 2C), γ-tubulin, ODF2, pericentrin, and GCP6 (not shown), which suggests that the apicosome represents either a group of vesicles or a microtubule-based organelle surrounding the ncMTOC that does not contain γ-tubulin (Zheng et al., 2020). Second, the apicosome did not show staining for markers of other subcellular organelles such as the Golgi apparatus (Glogin97), mitochondria (Tom20), and early/late endosomes (Rab5/7/11) (not shown). Third, the apicosome is not the HB, a specialized structure of the ER membrane and mitochondria, because (1) the HB contains mitochondria, but the apicosome does not show staining for the mitochondrial marker Tom20; (2) the HB localizes below the fonticulus, on the side opposite to the apicosome; (3) the HB is only present in OHCs, whereas the apicosome is present in both OHCs and IHCs; and (4) the HB is considerably larger than the apicosome, ∼3 μm in diameter (Lim, 1986; Mammano et al., 1999). Because methanol fixation denatures F-actin (and other proteins) and weakens the phalloidin staining that marks the hair-cell boundary, we used PFA fixation in subsequent experiments for clear visualization of the hair-cell boundary and apicosome itself.

We next examined the developmental profile of the apicosome in cochlear hair cells. No apicosome signal was detected in embryonic day (E) 16 hair cells (Fig. 3A), but by around E18, the apicosome was observed as a cluster of small particles or vesicles at the neural side of the peri-cuticular necklace (Fig. 3A, B); this presumably reflects the initial assembly of the apicosome. Notably, between P0 and P4, the apicosome descended from the peri-cuticular necklace to the perinuclear region of OHCs (Fig. 3A). This timing of apicosome translocation depended on the cell type and tonotopic gradient. First, the translocation occurred earlier in OHCs than IHCs within the same region of the organ of Corti: the apicosome descended in most OHCs by P5 but remained subapical in nearly all IHCs at this stage (Fig. 3A); the apicosome descended to the perinuclear region and became dispersed in all hair cells only after P7 (Fig. 3A). Second, the apicosome remained subapical in all the IHCs in the apex of the cochlea at P5 but showed subapical localization in <50% of the IHCs in the cochlear base (Fig. 3C); this suggests that apicosome translocation occurs earlier at the basal than apical turns of the cochlea. By contrast, the apicosome persisted in adult (P50) vestibular hair cells (Fig. S2), although it became smaller and more distant from the apical membrane compared to that in neonatal vestibular cells (Fig. 1B). Because kinocilia disappear after P10 in cochlear hair cells and persist in neonatal and adult vestibular cells, similar to the expression of the apicosome in cochlear and vestibular hair cells, we speculate that the presence of the apicosome is associated with kinocilium maintenance. In addition, kinocilia in OHCs appear to be resorbed earlier than those in IHC in mice and gerbils (Jia et al., 2009; Lim and Anniko, 1985), consistent with the timing of apicosome translocation and disappearance.

**Figure 3.**
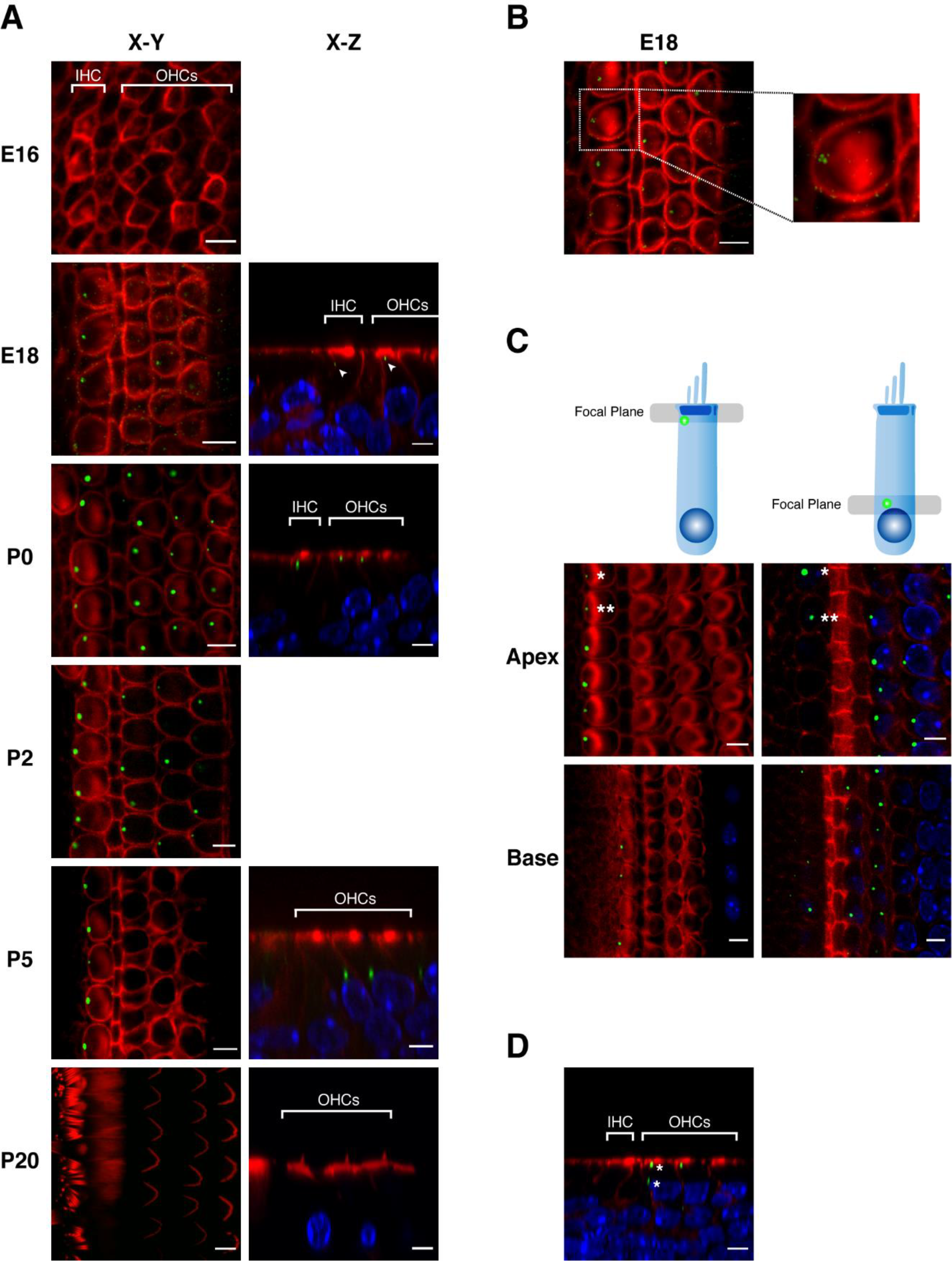
Apicosome developmental profile observed after PFA fixation. (**A**) X-Y scans of cochlear hair cells from WT mice at indicated ages (left column); some of their X-Z planes (right column) are also shown. Arrowheads: small particles or vesicles (presumably formed during apicosome assembly) detected in X-Z scans in both IHCs and OHCs at E18. (**B**) X-Y scan of E18 cochlear hair cells (left) and a magnification of one IHC (right). The apicosome (right), presumably still undergoing assembly, appears as a cluster of small particles or vesicles instead of as the single spot detected in more mature hair cells. The small particles also appear to surround cortical actin, presumably at the lateral cell surface. (**C**) Apicosome distribution in apical and basal turns of the cochlea from P5 WT mice. In upper schematic, gray bars indicate focal planes of images in lower panels, and positions of green dots represent typical locations of apicosomes at subapical or perinuclear regions. Panels in same row: images obtained at different focal planes from the same area; asterisks: appearance of 2 apicosomes at distinct focal planes in same cell. (**D**) X-Z scan of hair cells from P0 WT mouse. Asterisks: concurrent appearance of 2 apicosomes at subapical and perinuclear positions in same OHC. All panels: green, anti-FLRT1; red, phalloidin staining of F-actin; blue, DAPI staining of nucleus. Scale bars: 5 μm.

Interestingly, in a few hair cells, the apicosome was split and appeared concurrently at the subapical and perinuclear positions (Fig. 3C, D; Supplementary Video 1). By contrast, in most hair cells, the apicosome was found at either the subapical position or the perinuclear position as an entire entity and was seldom detected between the two positions (Fig. 3A). Thus, the apicosome might move from the apical position to the perinuclear position presumably in small steps, similar to the initial assembly (Fig. 3A, B), but this might occur too rapidly to be recorded in most hair cells. Alternatively, the apicosome might translocate as a single entity.

Sporadically, supernumerary IHCs was observed in the cochlea in WT mice (Figs. 4A and 1A); this might reflect minor fluctuations or errors in the control system of lateral inhibition during normal hair-cell development (Verpy et al., 2011) (Fig.2b in their study). The apicosome in these regions exhibited atypical features: (1) some of the hair cells contained more than one apicosome, and (2) the position of the apicosome was considerably outside its normal position (Figs. 4A and 1A). Thus, the abnormal appearance of the apicosome appears to be potentially associated with lateral inhibition. To further test this idea, we treated cochlear organ cultures with the γ-secretase inhibitor DAPT to disrupt Notch signaling pathway/lateral inhibition and generate supernumerary hair cell-like cells from trans-differentiation of supporting cells. Notably, the apicosome appeared in the supernumerary hair cells that are converted from supporting cells, further strengthening the notion that the apicosome is linked with hair-cell specification and differentiation (Fig. 4B). Moreover, the apicosome in the DAPT-induced supernumerary hair cells also show abnormal features similar with those in Fig. 4A (Fig. 4B), and these results bolster the association of atypical apicosome appearance with abnormal lateral inhibition. However, the apicosome did not appear to contain Delta-like ligand1 (DLL1) of Notch receptors, based on our immunostaining study with an antibody against DLL1 (not shown).

**Figure 4.**
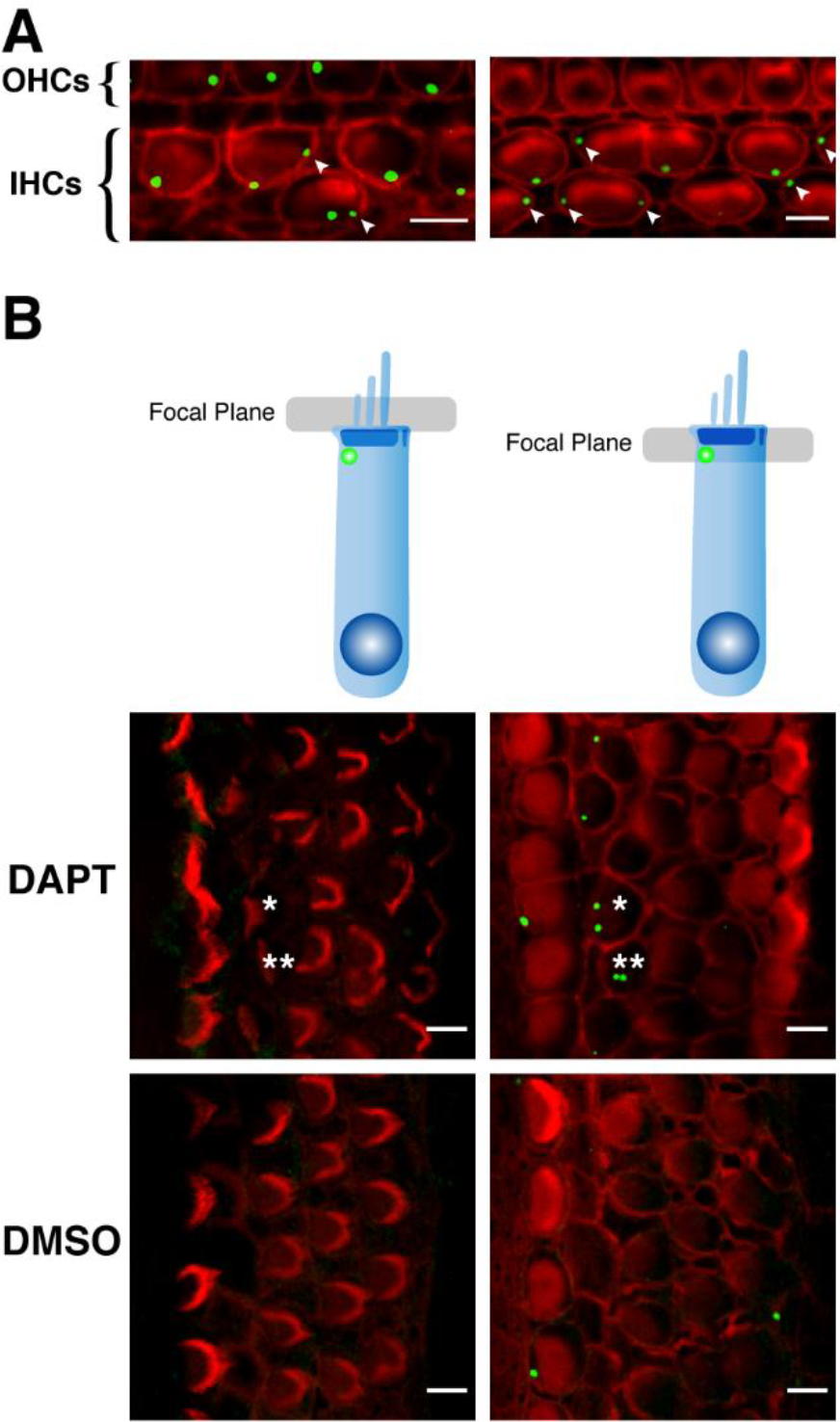
Apicosome localization in supernumerary HCs. (A) Apicosomes in supernumerary IHCs from P2 (left) and P5 (right) WT mice. Arrowheads: apicosomes exhibiting atypical localization—not at the neural side of hair cells. (**B**) Atypical localization of the apicosome is also seen in supernumerary HCs (P0 + 5 DIV) induced by DAPT treatment in apical-middle turns of WT cochlear cultures. The apicosomes already translocate and are not seen in the focal planes of these images in most of the cells. Asterisks: the same cell at different focal planes. Green: anti-FLRT1; red: phalloidin staining of F-actin. Scale bars: 5 μm.

Lastly, to gain additional insights into the biological nature of the apicosome, we examined whether the apicosome was affected when cochlear cultures were exposed to various reagents; we treated P2+1DIV organ of Corti cultures for 6 h with following reagents: BAPTA-AM or thapsigargin, for reducing or increasing intracellular Ca^2+^ levels; DHS, for blocking the MT channel (and hair-cell maturation); JW67, for inhibiting Wnt signaling; and EGF. However, the apicosome showed no change after each treatment and remained at the peri-cuticular necklace under the apical surfaces in all IHCs (Fig. 5).

**Figure 5.**
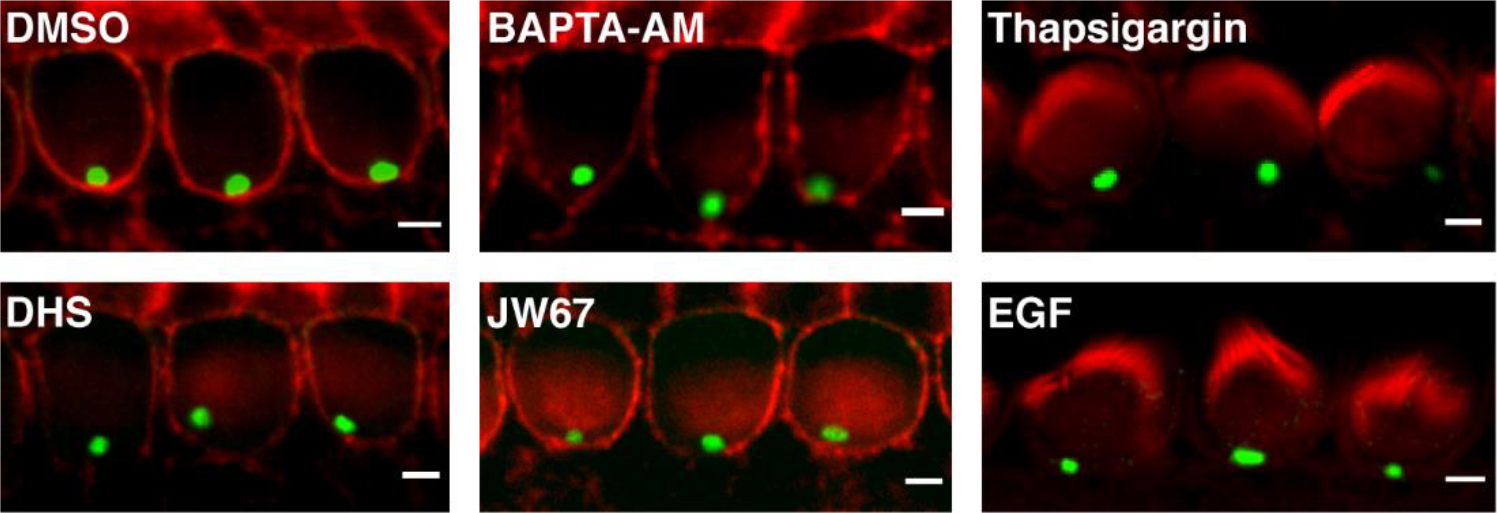
Apicosomes remain stable under various treatments. Apicosomes and F-actin were stained with anti-FLRT1 (green) and phalloidin (red), respectively, in P2+1DIV IHCs after treatment with indicated reagents for 6 h at 37°C. DMSO, solvent control. Scale bars: 2 μm.

## DISCUSSION

We identified an organelle exhibiting a unique localization, translocation, and developmental profile in both cochlear and vestibular hair cells, and have coined the name “apicosome” for this previously undescribed organelle (Fig. 6). Although the apicosome was fortuitously detected based on its recognition by two anti-FLRT1 antibodies, the organelle does not contain FLRT1, because anti-FLRT1 detected the apicosome even in *Flrt1*^*-/-*^ mice (Fig. 1). Moreover, FLRT1 itself does not appear to be involved in any aspect of hair-cell function or development: the results of ABR assays showed that hearing function was normal in 1-month-old *Flrt1*^*-/-*^ mice (Fig. S4).

**Figure 6.**
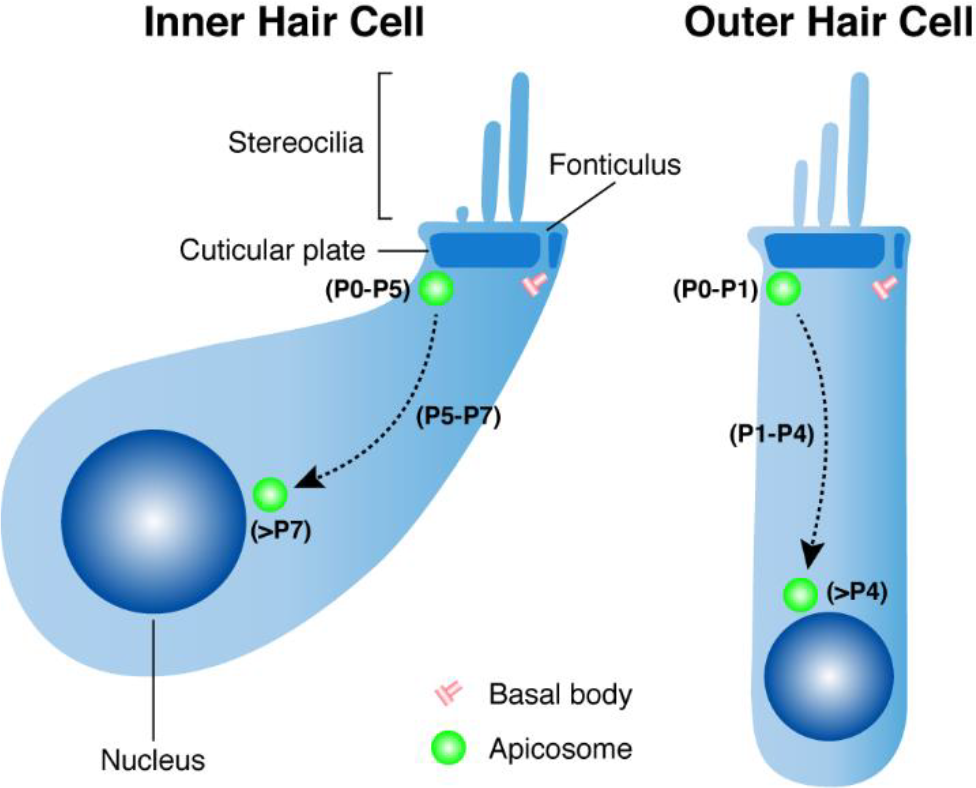
Schematic depicting apicosome translocation in IHCs and OHCs. Apicosomes (green) in IHCs remain at the neural side of the subapical region from P0 to P5, start to descend at P5, and all reach the perinuclear region after P7. Conversely, apicosomes in OHCs, although also assembled at E18 as in IHCs, remain at the subapical region for a shorter time (P0–P1) before descending, and all are localized at the perinuclear region after P4. Apicosomes in both IHCs and OHCs disintegrate after reaching the perinuclear region. Notably, at the tonotopic level, apicosome translocation (in both IHCs and OHCs) occurs earlier at the basal cochlea than at the apical cochlea. Stereocilia, cuticular plate, fonticulus, basal bodies, and nuclei are shown in both cell types.

Could the apicosome contain the FLRT1 homologs FLRT2 and FLRT3? FLRT2 and FLRT3 share high sequence similarity with FLRT1 (61% and 71%, respectively) (Lacy et al., 1999), and, more importantly, FLRT3 might be involved in hearing: an FLRT3 mutation (205C>A) was reported to be associated with hearing loss through the FGF signaling pathway (Miraoui et al., 2013). However, the apicosome was not recognized by anti-FLRT2 and anti-FLRT3 antibodies (Fig. S3) whose specificity was validated here (Fig. S3A, B; data not shown for anti-FLRT3). FLRT ectodomains contain metalloprotease cleavage sites and can be shed from FLRT-expressing cells (Yamagishi et al., 2011); thus, we reasoned that blocking the cleavage of the ectodomains of cell-surface FLRTs might alter apicosome size or dynamics if the apicosome acts as an intracellular pool of FLRTs. However, 10-h treatment of organ of Corti cultures (P0+2DIV) with metalloprotease inhibitors (20 μg/mL TAPI-0 and TAPI-1) did not affect apicosome localization or dynamics (not shown). To further identify candidate target proteins recognized by anti-FLRT1 in the apicosome, we performed BLAST searches by using the FLRT1 sequences aa 63–253, 50–150, and 63–150, the antigens of the two anti-FLRT1 antibodies (ab97825 and ab103839) and their overlapping region. However, all the BLAST searches yielded only FLRT1 proteins from various species. One possibility is that the two anti-FLRT1 antibodies recognize a secondary or tertiary structure rather than a primary sequence in the apicosome. Thus, mass spectrometry analysis of proteins immunoprecipitated using the anti-FLRT1 antibodies might represent an effective approach for addressing this matter. A precedent here can be found in the identification of PCDH15: in the tip link of hair cells, PCDH15 was first detected by an unidentified antibody, and the molecular identity of PCDH15 was only revealed later by using mass spectrometry (Ahmed et al., 2006).

The translocation trajectory of the apicosome before complete disappearance was found to be highly characteristic and consistent—from the subapical location at the neural side to the perinuclear position. This is reminiscent of the translocation of certain latent gene regulatory proteins such as Notch, β-catenin, and Smads; these molecules traffic from the cell surface or the cytoplasm into the nucleus and function in lateral inhibition, planar cell polarity signaling, or pattern formation. However, these signaling molecules are all translocated as soluble proteins and require no membranous vesicle such as the apicosome. Interestingly, apicosome translocation itself might be mediated by microtubules, because our results (Fig. 2) suggested that the apicosome represents either a group of vesicles or a microtubule-based ncMTOC-surrounding organelle lacking γ-tubulin. Microtubules are known to be present throughout the cell body in both OHCs and IHCs, and some of these microtubes are concentrated in the tubulo-vesicular track (Furness et al., 1990) and could therefore serve as the track for apicosome translocation. Another potential track is the SO. Although the precise composition of the SO is unclear, it contains α2-spectrin, and the SO is also present throughout the cell body (Vranceanu et al., 2012). However, the SO is present in vestibular and cochlear IHCs but not OHCs (Vranceanu et al., 2012), whereas the apicosome is present in both IHCs and OHCs.

In summary, we serendipitously identified a previously unrecognized hair-cell-specific organelle, the apicosome; the organelle shows a distinctive localization, translocation, and developmental profile and is potentially associated with kinocilium maintenance and lateral inhibition. The itinerant nature and transient appearance of the apicosome during development might explain why the organelle remained unnoticed over the past several decades of research on hair cells. The critical question to address next is the molecular composition of the apicosome, including the protein recognized by anti-FLRT1; this will be essential for elucidating the function, regulation, and translocation of the apicosome. Nevertheless, our work lays the foundation for further characterization of this unique structure potentially linked to hair-cell development and function.

## ACKNOWLEDGMENTS

We thank Prof. Yusong Guo and Dr. Pengfei Liu for helpful discussion and suggestions. This work was supported by Hong Kong RGC GRF16102417 and GRF16100218, NSFC-RGC Joint-Research Scheme N_HKUST614/18, Shenzhen Basic Research Scheme (JCYJ20170818114328332), SMSEGL20SC01-K, and in part by the Innovation and Technology Commission (ITCPD/17-9).

## AUTHOR CONTRIBUTIONS

Conceptualization: X.L., X.Y., and P.H.; methodology: X.L., L.J., D.Z.H., and R.Z.Q.; formal analysis: X.L. and P.H.; investigation: X.L., Q.Z., X.Y., W.C., Y.Z., and W.F.; resources: L.J., D.Z.H. and R.Z.Q; writing-original and draft: X.L. and P.H.; writing-review and editing: P.H., with contributions from all the authors; visualization: X.L.; supervision: P.H.; funding acquisition: P.H.

