## Supplementary Materials for "Apicosome: newly identified cell-type-specific organelle in mouse cochlear and vestibular hair cells"

### SUPPLEMENTARY VIDEO AND FIGURES

**Supplementary Video 1. 3D reconstruction of apicosomes in cochlear OHCs and IHCs in P4 WT mouse.** The 3D video was reconstructed using images of 20 consecutive X-Y sections (0.5-µm interval) acquired using confocal microscopy. Apicosomes (green) in IHCs display mostly homogenous size, shape, and localization; apicosomes in some of the OHCs have descended to the perinuclear position. Red, phalloidin staining of actin (tuned down intentionally for clear visualization of apicosomes).

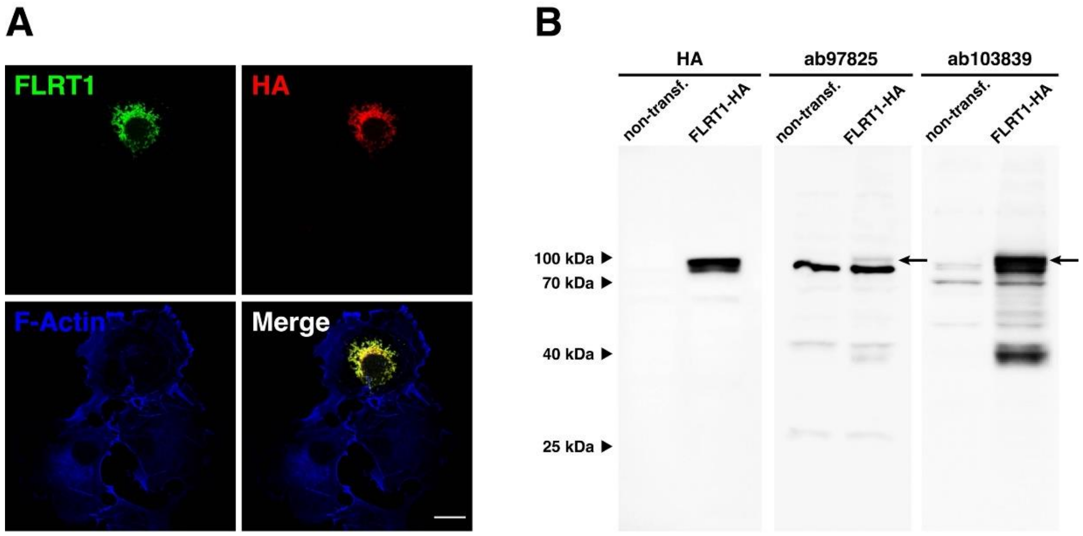


**Figure S1. Validation of anti-FLRT1 specificity.** (**A**) COS7 cells transfected with C-terminal HA-tagged FLRT1 (FLRT1-HA) were immunostained with anti-FLRT1 (ab97825, Abcam) and anti-HA. Scale bar: 20 μm. (**B**) HEK293T cells were not transfected (non-transf.) or transfected with FLRT1-HA and then immunoblotted with two distinct anti-FLRT1 antibodies (ab97825 and ab103839; 1:1000) and anti-HA (1:1000). Arrows: FLRT1 bands recognized by anti-FLRT1 antibodies.

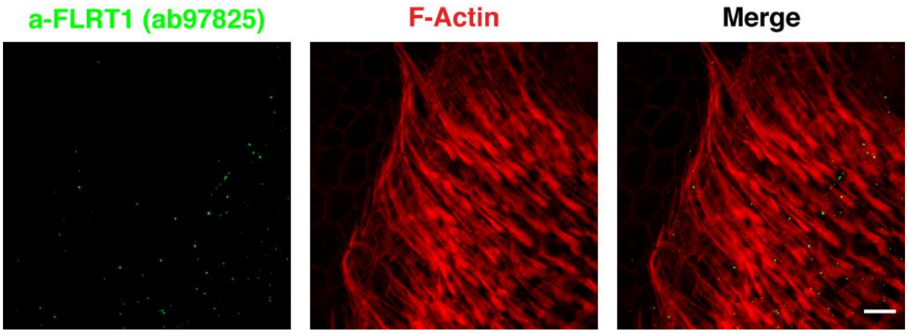


**Figure S2. Apicosomes in vestibular hair cells from adult WT mice.** Apicosomes were observed in P50 vestibular hair cells but absent in supporting cells (left region, weak appearance of cell-boundaries), and shown is a stack of images acquired within a depth of 15 μm. Scale bar: 10 μm.

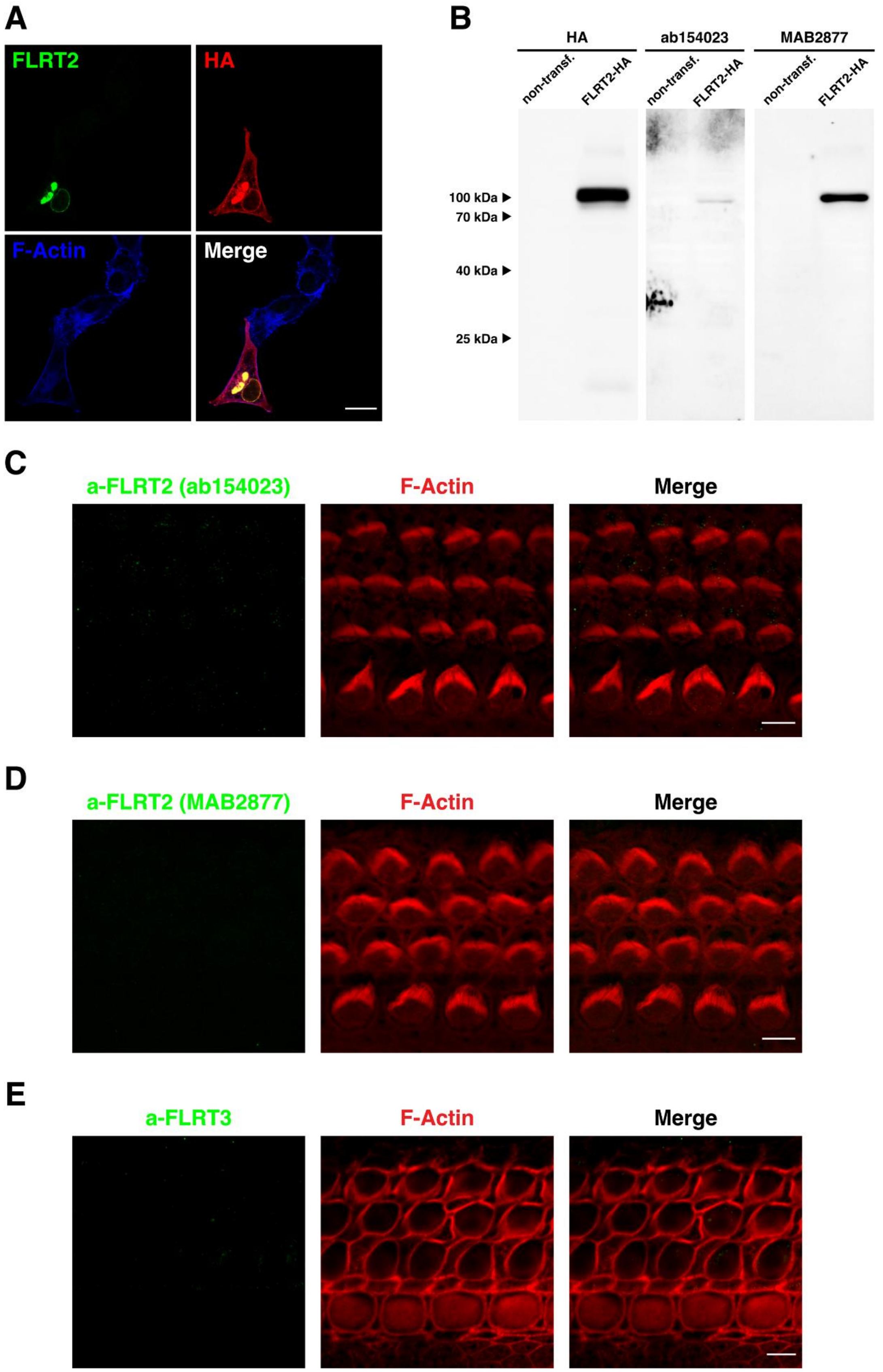


**Figure S3. Validation of anti-FLRT2 specificity and immunostaining of cochlear hair cells with anti-FLRT2 and anti-FLRT3.** (**A**) COS7 cells transfected with C-terminal HA-tagged FLRT2 (FLRT2-HA) were immunostained with anti-FLRT2 (ab154023, Abcam) and anti-HA. Anti-FLRT2 and anti-HA signals inside the cell largely overlap. Notably, the anti-FLRT2 signal is not detected on the cell surface; one simple explanation is that the anti-FLRT2 epitope in cell-surface FLRT2 is less accessible than the intracellular epitope. (**B**) Western blotting of non-transfected (non-transf.) and FLRT2-HA-transfected HEK293T cells with two anti-FLRT2 antibodies (ab154023 and MAB2877; 1:1000) and anti-HA (1:1000). (**C-D**) Anti-FLRT2 antibodies ab154023 and MAB2877 fail to stain apicosomes in apical-middle cochleae from P2 (C) and P3 (D) WT mice, respectively. (**E**) Anti-FLRT3 fails to recognize apicosomes in cochlear hair cells from P6 WT mice. Phalloidin staining shows F-actin (red). Scale bars: 20 μm (A), 5 μm (C-E).

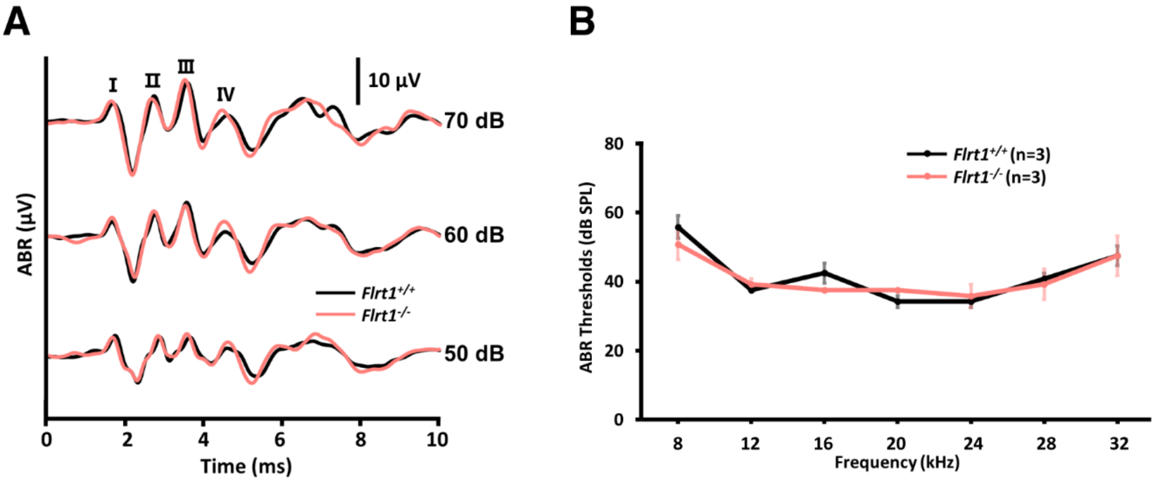


**Figure S4. Hearing function of *Flrt1*$^{-/-}$ mice.** Representative ABR traces elicited by click stimuli (**A**) and ABR thresholds in response to pure tones (**B**) in P28–P32 *Flrt1*$^{+/+}$ (n=3) and *Flrt1*$^{-/-}$ mice (n=3). No hearing impairment was found in *Flrt1*$^{-/-}$ mice. Data in (B) are expressed as means ± SEM; n, number of independent biologic replicates. Two-tailed Student's *t* test was used for statistical analysis, and $P \geq 0.05$ was considered statistically insignificant.
